# The third coiled coil domain of Atg11 is required for shaping mitophagy initiation sites

**DOI:** 10.1101/2020.06.29.178780

**Authors:** Hannah K. Margolis, Sarah Katzenell, Kelsie A. Leary, Michael J. Ragusa

## Abstract

Selective autophagy is the capture of specific cytosolic contents in double membrane vesicles that subsequently fuse with the vacuole or lysosome, thereby delivering cargo for degradation. Selective autophagy receptors (SARs) mark the cargo for degradation and, in yeast, recruit Atg11, the scaffolding protein for selective autophagy initiation. The mitochondrial protein Atg32 is the yeast SAR that mediates mitophagy, the selective autophagic capture of mitochondria. Atg11- Atg32 interactions concentrate Atg32 into puncta that are thought to represent sites of mitophagy initiation. However, it is unclear how Atg11 concentrates Atg32 to generate mitophagy initiation sites. We show here that the coiled coil 3 (CC3) domain of Atg11 is required for concentrating Atg32 into puncta. We determined the structure of the majority of the CC3, demonstrating that the CC3 forms a parallel homodimer whose dimer interface is formed by a small number of hydrophobic residues. We further show that the CC3 dimerization interface is required for shaping Atg32 into functional mitophagy initiation sites and for delivery of mitochondria to the vacuole. Our findings suggest that Atg11 self-interactions help concentrate SARs as a necessary precondition for cargo capture.

## Introduction

Autophagy, a catabolic cellular process that is conserved from yeast to mammals, leads to the capture of cytosolic material in double membrane vesicles, termed autophagosomes (Dikic and Elazar, 2018; Wen and Klionsky, 2016). Once autophagosome biogenesis is completed, these vesicles fuse with the vacuole in yeast and plants, or lysosomes in animal cells, leading to the degradation of the captured cytosolic contents. The capture of autophagic cargos can occur by a non-selective or a selective mechanism. In selective autophagy, cargos, including aggregated proteins, pathogens, mitochondria, endoplasmic reticulum (ER), and other organelles, are marked for degradation by selective autophagy receptors (SARs) (Gatica et al., 2018; Zaffagnini and Martens, 2016). In yeast, SARs recruit Atg11, a cytoplasmic protein that serves as the scaffold for selective autophagy initiation. Once bound to SARs, Atg11 recruits other autophagy components to mediate autophagosome biogenesis (Zientara-Rytter and Subramani, 2020).

Atg32 is the yeast SAR that mediates mitophagy, the selective autophagy of mitochondria (Kanki et al., 2009b; Okamoto et al., 2009; Pickles et al., 2018). This single-pass transmembrane protein is localized to the outer mitochondrial membrane (OMM). In mitophagy-inducing conditions, Atg32 is activated by a series of post-translational modifications leading to the recruitment of Atg11 (Aoki et al., 2011; Farré et al., 2013; Kanki et al., 2013; Wang et al., 2013). Atg11 binding to Atg32 triggers the formation of Atg32 puncta on the surface of mitochondria (Furukawa et al., 2018). These puncta initially colocalize with Atg11, then with the autophagosomal marker Atg8, and finally are delivered to the vacuole (Furukawa et al., 2018). Thus, Atg32 puncta on the surface of mitochondria have been proposed to represent the site of mitophagy initiation (Yamashita and Kanki, 2017). While it has been established that Atg11 is required for the formation of Atg32 puncta, its role in this process is not yet known.

Atg11 is a 1178 amino acid cytoplasmic scaffolding protein that binds to all SARs in yeast to initiate selective autophagy (Yorimitsu and Klionsky, 2005). No structures have been determined for any region of Atg11, but secondary structure predictions suggest that it contains 4 coiled coil (CC) domains (CC1-4, Figure 1A). The N-terminal region of Atg11 spans residues 1-576 and contains the first two coiled coil domains (CC1 and CC2). This region binds the core autophagy factors Atg1 (homologue of the mammalian ULK1), Atg13, and Atg9 (Chang and Huang, 2007; Kamber et al., 2015; Matscheko et al., 2019; Torggler et al., 2016). The C-terminal region spans residues 860-1178 and includes CC4 (Yorimitsu and Klionsky, 2005; Zientara-Rytter and Subramani, 2020). This region binds SARs, including Atg32 and Atg19 (Aoki et al., 2011; Yorimitsu and Klionsky, 2005). Two crystal structures were recently determined for the C-terminal region of FIP200, the proposed mammalian analog of Atg11. These structures revealed the presence of a CLAW domain at the extreme C-terminus of FIP200, which like the CTD of Atg11, is responsible for binding SARs (Turco et al., 2019). Based on sequence homology to FIP200 it is likely that Atg11 contains a similarly structured CLAW domain at its C-terminus for SAR binding.

**Figure 1.**
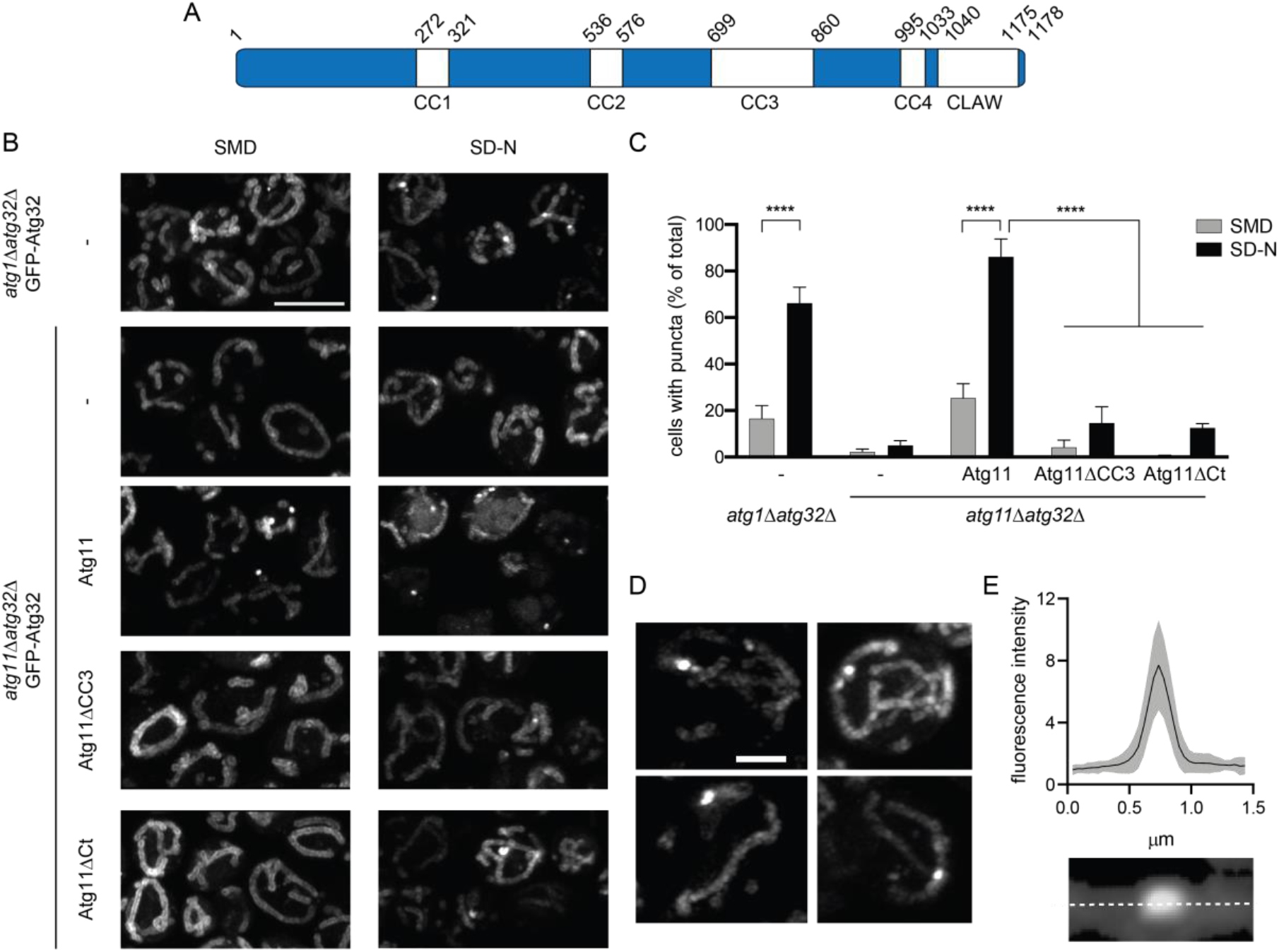
Atg11CC3 is required for the formation of Atg32 puncta. **A.** A schematic of the domain architecture of Atg11 with the amino acid number for the start and end of each domain listed. **B.** Representative images showing GFP-Atg32 driven by the CUP1 promoter with 0.5 μM copper in either *atg1Δatg32Δ* or *atg11Δatg32Δ* cells. Where noted, *atg11Δatg32Δ* cells were also expressing the indicated Atg11 constructs. Cells were grown in SMD, then transferred to SD-N for 1 hr. Scale bar: 5μm. **C.** Quantification of the percent of cells containing puncta from B. Three independent repeats were performed with a total of >1000 cells analyzed per group. Significance was determined using a two-way ANOVA with Sidak’s multiple comparison test. **** p<0.0001. **D.** Representative images showing the variation in shape of GFP-Atg32 puncta in *atg1Δatg32Δ* cells incubated in SD-N for 1h. Scale bar: 2μm. **E.** MFI of GFP-Atg32 puncta and surrounding mitochondria. GFP fluorescence was measured along the dotted line and normalized to baseline fluorescence of Atg32 in the surrounding mitochondria. Measurements were made for 50 puncta imaged in 3 separate experiments. Black line – mean MFI; gray border – SD.

The longest of the Atg11 coiled coil domains is the CC3, which is predicted to span residues 699-860, bridging the N and C terminal regions of Atg11. Very little is known about the role that the CC3 plays in Atg11 function. Given its location between the N-terminal and C-terminal regions of Atg11 it may simply be a spacer separating the core autophagy machinery from the cargo. However, there is some evidence to suggest that its role in selective autophagy may be more complicated. The CC3 has been proposed to dimerize, although the role of this potential dimerization has not been explored (Yorimitsu and Klionsky, 2005). The CC3 has no known binding partners apart from Atg11, including any of the core autophagy machinery. Nonetheless, deletion of the entire CC3 prevents recruitment of the core autophagy machinery to the phagophore assembly site (PAS) (Yorimitsu and Klionsky, 2005). The PAS serves as the initiation site for cytoplasm to vacuole targeting (Cvt), a form of selective autophagy in yeast that delivers proteases to the vacuole (Lynch-Day and Klionsky, 2010). Thus, it seems plausible that the CC3 plays an important structural role during the initiation of selective autophagy. Given its potential structural role in PAS formation, we wondered if the CC3 might also play a role in the formation of mitophagy initiation sites.

## Results

### Airyscan Imaging of Atg32 puncta

We used Airyscan confocal microscopy to monitor the formation of GFP-Atg32 puncta as a marker of mitophagy initiation sites. Airyscan confocal microscopy achieves a resolution of 140 nm laterally and 400 nm axially and provides high sensitivity and a high signal-to-noise ratio. This enabled us to discern small differences in the structure and shape of mitophagy initiation sites upon perturbation of Atg11. Atg32 puncta have been shown to accumulate on mitochondria when *atg1* is deleted in yeast (Furukawa et al., 2018). To confirm this result, GFP-Atg32 was expressed in *atg1Δatg32Δ* cells that were grown in nitrogen rich media (SMD) and transferred to media lacking nitrogen sources for 1 hr (SD-N). In agreement with previous findings, we observed that Atg32 was restructured from a diffuse mitochondrial pattern into puncta that were retained on mitochondria (Figure 1B and C) (Furukawa et al., 2018; Mao et al., 2011). We measured the diameter and brightness of Atg32 puncta and found that most were circular or slightly ovular in shape and were 0.39 ± 0.12 μm in diameter (Figure 1D and E). The mean fluorescence intensity (MFI) of Atg32 puncta was 7.7 times brighter than that of the surrounding mitochondria (Figure 1E), demonstrating that Atg32 puncta represent sites where Atg32 is highly concentrated.

To confirm the requirement of Atg11 for Atg32 puncta formation, we quantified puncta abundance in *atg11Δatg32Δ* cells. In contrast to *atg1Δatg32Δ* cells, *atg11Δatg32Δ* cells had essentially no Atg32 puncta, even after transfer to SD-N (Figure 1B and C). These data confirm that Atg11 is required for the formation of Atg32 puncta. We found similar results when Atg32 was expressed at either lower or higher levels on mitochondria (Supplemental Figure S1A), suggesting that the concentration of Atg32 in the OMM does not affect the formation of Atg32 puncta. We also found similar results using mCherry-Atg32 (Supplemental Figure S1B and C), suggesting that the fluorophore used did not affect Atg32 puncta formation. We concluded that, following a mitophagy-inducing stimulus, Atg32 is reorganized from a diffuse mitochondrial localization into distinct foci in an Atg11-dependent manner.

### Atg11CC3 is required for Atg32 puncta formation

Since there is currently no established role for the CC3 in mitophagy initiation, we wondered if it was required for the reorganization of Atg32 into puncta. To test this, we generated a set of Atg11 constructs that were driven by the endogenous Atg11 promoter, including full-length Atg11 (Atg11), Atg11 lacking residues 699-860 (Atg11ΔCC3), and Atg11 lacking residues 965-1178 (Atg11ΔCt), which is deficient in Atg32 binding (Aoki et al., 2011). We co-expressed these constructs with GFP-Atg32 in *atg11Δatg32Δ* cells and monitored the formation of Atg32 puncta. As expected, expression of Atg11 restored the formation of Atg32 puncta (Figure 1B) and GFP- Atg32 was delivered to the vacuole in many cells, suggesting that mitophagy was functional in those cells. In contrast, Atg11ΔCt did not restore Atg32 puncta formation or vacuolar GFP, likely due to the loss of Atg32 binding (Figure 1B and C). Surprisingly, Atg11ΔCC3 showed a comparable reduction in Atg32 puncta formation. Moreover, there was no evidence of GFP delivery to the vacuole in cells expressing Atg11ΔCC3 (Figure 1B and C). This suggests that mitophagy initiation sites were unable to form in the absence of the Atg11CC3 and that mitophagy is abolished as a result.

One possible explanation for the loss of Atg32 puncta in cells expressing Atg11ΔCC3 is that deletion of the CC3 leads to a loss of Atg11 binding to Atg32. To test this, we expressed GFP- Atg32 with FLAG-Atg11, FLAG-Atg11ΔCC3 or FLAG-Atg11ΔCt and performed a co-immunoprecipitation assay. We found that Atg11ΔCC3 was still able to pull down Atg32, although there was a reduction in binding compared to full-length Atg11. In contrast, Atg11ΔCt was unable to pull down any Atg32, consistent with the role of the C-terminus of Atg11 in SAR binding (Figure 2A). It has been proposed that SAR clustering promotes cargo capture by increasing the avidity for binding FIP200 and other autophagy proteins (Wurzer et al., 2015; Zaffagnini and Martens, 2016) (Turco et al., 2019). Since Atg11ΔCC3 cannot cluster Atg32, it may have a reduced avidity for Atg32, which would explain why we observed a decrease in Atg32 binding to Atg11ΔCC3 as compared to full-length Atg11. Thus, it seems that the CC3 enhances Atg11 binding to Atg32 but is not absolutely required for Atg32 binding.

**Figure 2.**
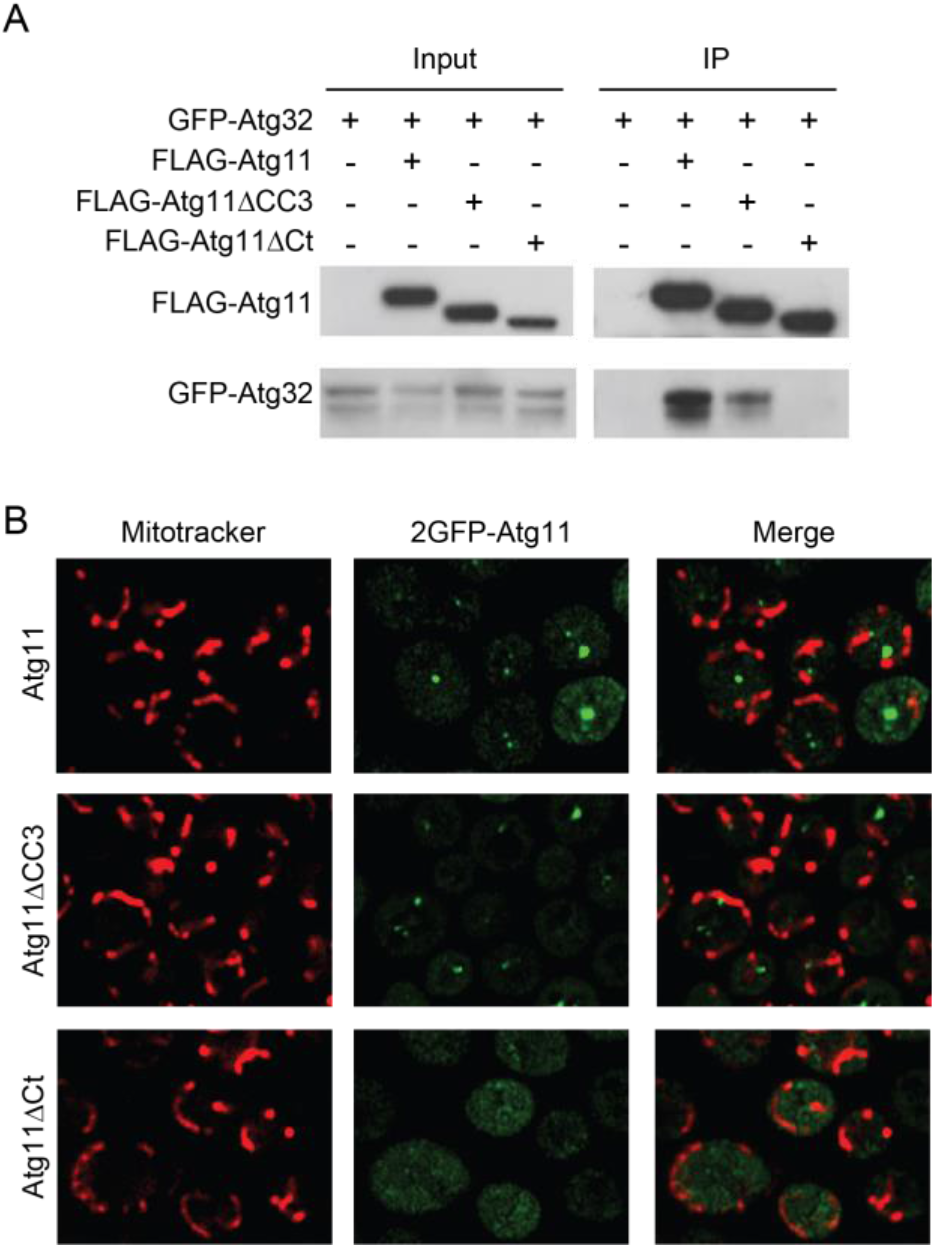
Atg11ΔCC3 is still recruited to mitochondria during mitophagy induction. **A.** GFP-Atg32 and FLAG-Atg11, FLAG-Atg11ΔCC3, or FLAG-Atg11ΔCt were expressed in *atg11Δatg32Δ* cells. Cells were grown in SMD and then transferred to SD-N media for 30 minutes. Lysate was immunoprecipitated with anti-FLAG resin and probed with anti-FLAG and anti-GFP antibodies. Experiments were performed in triplicate; representative blots are shown. **B.** Representative images of *atg11Δ* cells expressing 2GFP-Atg11 constructs. Cells were grown in SMD and then transferred to SD-N media for 1 hr, then stained with MitoTracker Red and imaged. Single plane images are shown.

Due to the reduction in binding between Atg32 and Atg11ΔCC3, we wanted to determine if removing the CC3 affected the recruitment of Atg11 to mitochondria. To test this, we expressed Atg11 constructs tagged with two tandem copies of GFP in *atg11Δ* cells. We found that 2GFP- Atg11 and 2GFP-Atg11ΔCC3 were both recruited to the mitochondrial surface following incubation in SD-N (Figure 2B). In contrast, 2GFP-Atg11ΔCt was not recruited to mitochondria (Figure 2B). These findings support the interpretation that, unlike the Atg11Ct, Atg11CC3 is not strictly required for Atg11 recruitment to the mitochondria or binding to Atg32. Thus, the near complete loss of Atg32 puncta formation and vacuolar delivery in cells expressing Atg11ΔCC3 must be attributable to some other function performed by the CC3.

### Atg11CC3 is a parallel coiled coil dimer

To explore how the CC3 might function in the formation of mitophagy initiation sites, we crystallized and determined the structure of Atg11 residues 699-800 (Atg11_699-800_) (Figure 3A and Table 1), which includes the majority of the CC3. The structure shows that this region forms a parallel coiled coil dimer. The asymmetric unit contained a single copy of Atg11_699-800_ and the crystallographic 2-fold symmetry mate contained the dimeric partner (Figure 3A). Electron density was observed for residues 699-784, leaving residues 785-800 unbuilt in the structure. Surprisingly, a large portion of the residues in the structure contained poor electron density and had high B-factors, suggesting that these regions of Atg11 are more dynamic than the central region of the coiled coil (Supplemental Figure S2).

**Figure 3.**
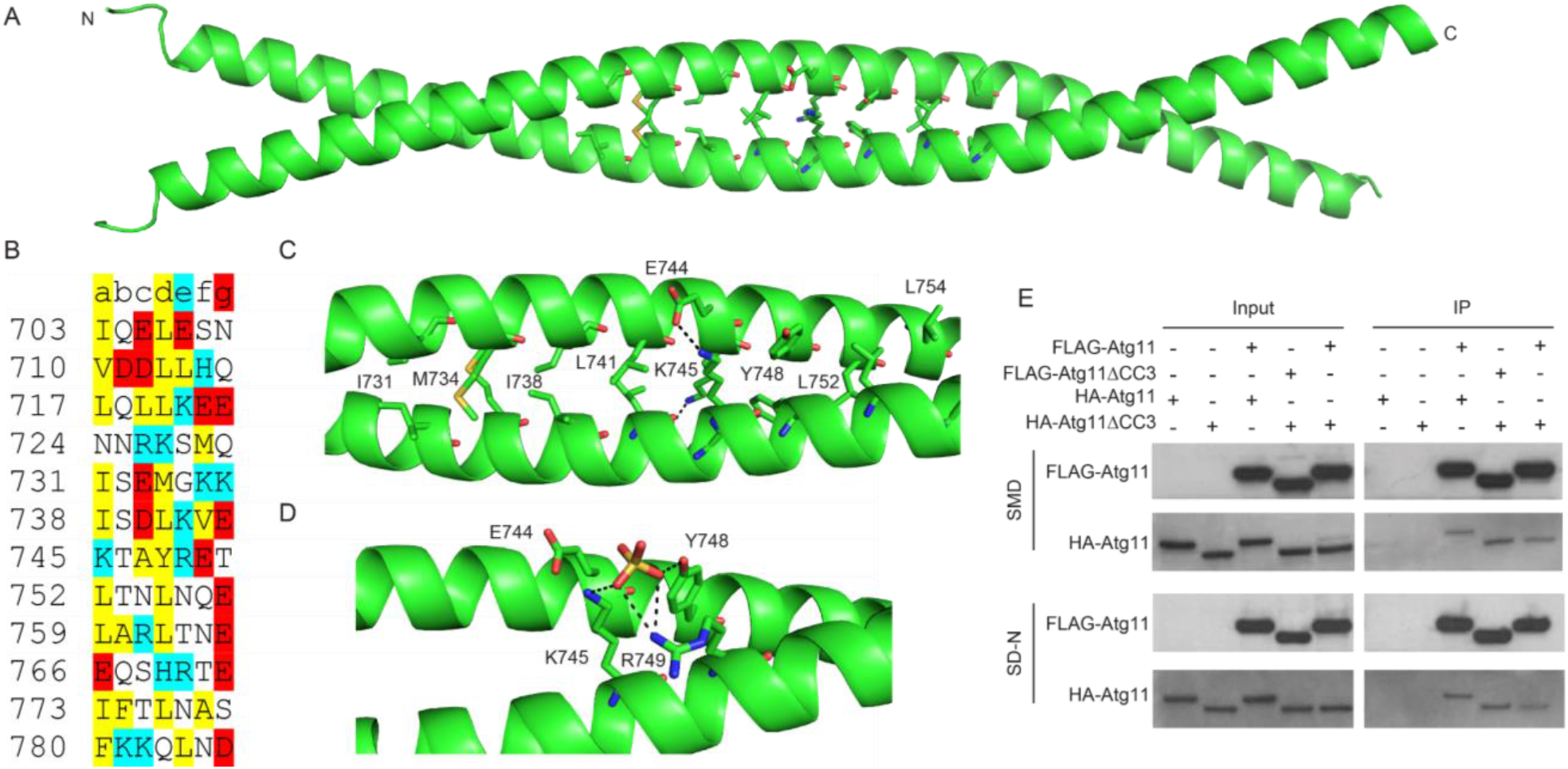
Crystal structure of Atg11_699-800_. **A.** Crystal structure of Atg11_699-800_ shown as a ribbon diagram. Side chains are shown for residues at the dimer interface. The N and C termini are labeled. **B**. *S. cerevisiae* Atg11 sequence from 703 to 786 organized by heptad repeat. Hydrophobic residues, positively charged and negatively charged residues are colored yellow, blue and red, respectively. For an ideal heptad repeat, hydrophobic residues should be in positions a and d while charged residues are in positions e and g. **C.** Close up view of the central region of the coiled coil dimer interface. Residues in A are shown as stick representations and labeled. Electrostatic interactions are shown as dashed lines. **D.** Residues involved in coordinating a sulfate ion are shown. Polar contacts are shown as dashed lines. **E.** FLAG-Atg11 or FLAG- Atg11ΔCC3 and HA-Atg11 or HA-Atg11ΔCC3 were expressed in *atg11Δ* cells. Cells were grown in SMD and then transferred to SD-N media for 30 minutes. Lysate was immunoprecipitated with anti-FLAG resin and probed with anti-FLAG and anti-HA antibodies. Experiments were performed in triplicate; representative blots are shown.

**Table 1 -.**
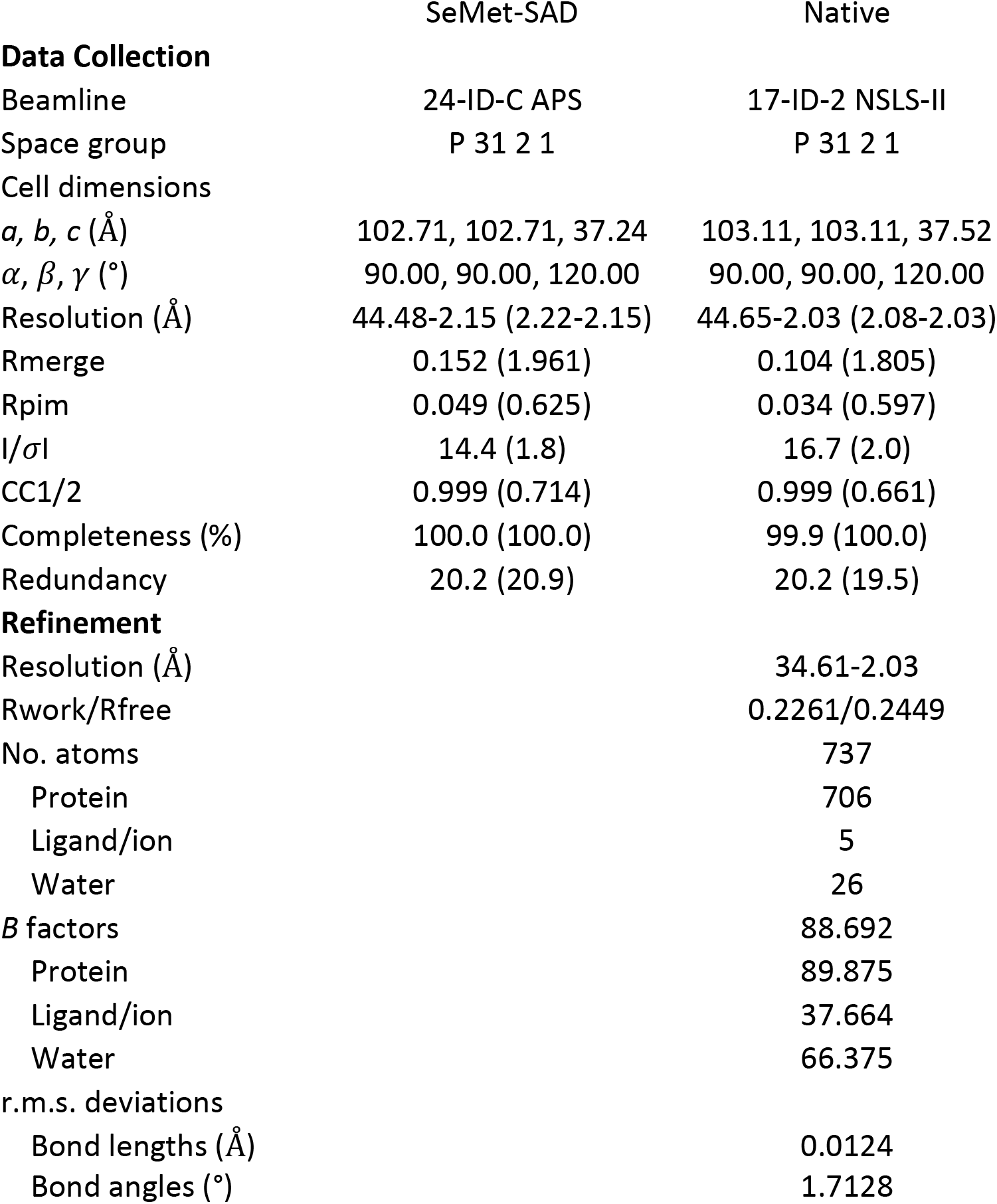
X-ray data collection and refinement statistics

|  | SeMet-SAD | Native |
| --- | --- | --- |
| **Data Collection** |  |  |
| Beamline | 24-ID-C APS | 17-ID-2 NSLS-II |
| Space group | P 31 2 1 | P 31 2 1 |
| Cell dimensions |  |  |
| *a*, *b*, *c* (Å) | 102.71, 102.71, 37.24 | 103.11, 103.11, 37.52 |
| *α*, *β*, *γ* (°) | 90.00, 90.00, 120.00 | 90.00, 90.00, 120.00 |
| Resolution (Å) | 44.48-2.15 (2.22-2.15) | 44.65-2.03 (2.08-2.03) |
| Rmerge | 0.152 (1.961) | 0.104 (1.805) |
| Rpim | 0.049 (0.625) | 0.034 (0.597) |
| *I*/*σ**I* | 14.4 (1.8) | 16.7 (2.0) |
| CC1/2 | 0.999 (0.714) | 0.999 (0.661) |
| Completeness (%) | 100.0 (100.0) | 99.9 (100.0) |
| Redundancy | 20.2 (20.9) | 20.2 (19.5) |
| **Refinement** |  |  |
| Resolution (Å) |  | 34.61-2.03 |
| Rwork/Rfree |  | 0.2261/0.2449 |
| No. atoms |  | 737 |
| Protein |  | 706 |
| Ligand/ion |  | 5 |
| Water |  | 26 |
| *B* factors |  | 88.692 |
| Protein |  | 89.875 |
| Ligand/ion |  | 37.664 |
| Water |  | 66.375 |
| r.m.s. deviations |  |  |
| Bond lengths (Å) |  | 0.0124 |
| Bond angles (°) |  | 1.7128 |

The region of the structure containing the clearest electron density and the lowest B-factors (≤ 60) was for residues 730-760 (Figure 3B and C and Supplemental Figure S2). Within this region, 6 hydrophobic amino acids (Ile731, Met734, Ile738, Leu741, Leu752, Leu755) sit at the interface between the two copies of the CC3. Coiled coil domains consist of heptad repeats that are defined by repeating sequences of 7 amino acids, labeled a through g. Most commonly, positions a and d are occupied by hydrophobic residues that pack in the center of the helices, while positions e and g are occupied by charged residues that form electrostatic interactions on the outside surface the helical bundle. The 6 hydrophobic residues in the 730-760 region do indeed conform to a standard heptad repeat (Figure 3B). The dimer interface is further supported by electrostatic interactions between Lys745 and Glu744. Together, these 8 residues appear to be the most important residues for supporting the coiled coil dimer interface. This central region of strong heptad repeats is flanked on either side by sequences (724-730 and 766-772) that lack hydrophobic residues in either position a or d, potentially contributing to the increased flexibility that we observed in the crystal structure (Figure 3B).

We also observed a sulfate ion packed between two coiled coil dimers of Atg11_699-800_ (Figure 3D). The packing of this sulfate ion is coordinated by two highly conserved residues, Lys745 and Arg749. Notably, Arg749 is located away from the coiled coil interface, suggesting that it may not be directly involved in CC3 dimerization. Since sulfate ions can dock at functionally important sites, including potential protein-protein interaction sites, these conserved residues may contribute to the function of the CC3 in mitophagy initiation. Taken together, our structural data shows that the 699-800 region of Atg11 forms a dimeric coiled coil, supported in its central region by eight interfacing residues that make up a stretch of clear heptad repeats.

### Atg11CC3 is not required for the overall dimerization of Atg11

Atg11 is known to dimerize *in vivo*, and recent studies on recombinantly expressed and purified Atg11 indicate that both the N- and C-terminal fragments are capable of dimerization (Suzuki and Noda, 2018; Yorimitsu and Klionsky, 2005). However, it is not clear whether the CC3 is required for Atg11 dimerization *in vivo*. To test this, we expressed FLAG-Atg11 or FLAG-Atg11ΔCC3 with HA-Atg11 or HA-Atg11ΔCC3 and performed a co-immunoprecipitation assay. We found that Atg11 co-immunoprecipitated in both SMD and SD-N media, confirming its capacity to self-associate independent of nutrient availability (Figure 3E). To our surprise, loss of CC3 did not diminish Atg11 co-immunoprecipitation (Figure 3E), even when both Atg11 constructs were lacking the CC3. This suggests that the CC3 is entirely dispensable for the overall dimerization of Atg11, even though the CC3 is itself a dimer.

### Atg11CC3 promotes the organization of mitophagy initiation sites

To test the importance of the CC3 dimerization interface for Atg32 puncta formation, we generated four Atg11 constructs: a deletion of the crystallized region (Atg11Δ699-800), a deletion of the well-ordered region (Atg11Δ731-765), a mutation of the 6 hydrophobic amino acids that form the dimer interface (Atg11Φ6A) to Ala, and a double mutation, Lys745Ser and Arg749Ser (Atg11-2Ser), to disrupt the sulfate docking site. We transformed these constructs along with GFP-Atg32 into *atg11Δatg32Δ* cells and monitored the formation of Atg32 puncta following SD-N treatment (Figure 4A and B). We found that cells expressing Atg11Δ699-800, Atg11Δ731-765, or Atg11Φ6A had significantly fewer Atg32 puncta than cells expressing full-length Atg11 and almost none of these cells had vacuolar GFP (Fig. 4A and B). This striking loss of function is particularly noteworthy in the Atg11Φ6A mutant, where there is no truncation and only a disruption of the CC3 dimerization interface.

**Figure 4.**
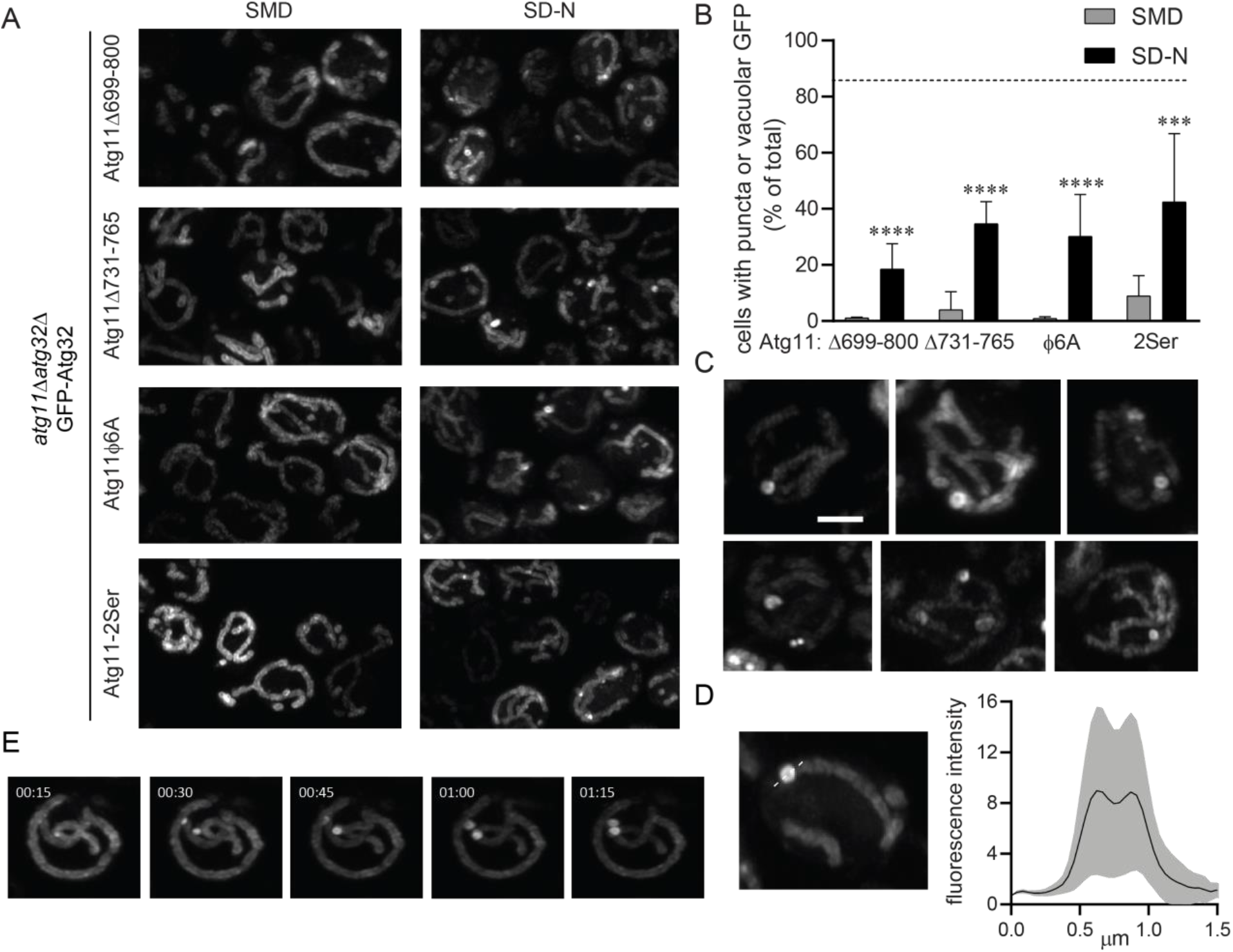
Coiled coil dimer interface mutants in the Atg11CC3 disrupt the formation of Atg32 puncta. **A.** Representative images of *atg11Δatg32Δ* cells expressing GFP-Atg32 and the indicated Atg11 constructs. **B** Quantification of the percent of cells containing puncta or vacuolar GFP from A. Three independent repeats were performed with a total of >1000 cells analyzed per group. Significance was determined using a two-way ANOVA with Sidak’s multiple comparison test. **** p<0.0001 *** p<0.001 and * p<0.05 **C.** Representative images showing the variation in crescent and ring-shaped GFP-Atg32 puncta in *atg11Δatg32Δ* cells expressing Atg11Δ731-765 and incubated in SD-N for 1 hr. Scale bar: 2μm. **D.** MFI of GFP-Atg32 rings and surrounding mitochondria shown in C. GFP fluorescence was measured along the dotted line and normalized to baseline fluorescence of Atg32 in the surrounding mitochondria. Measurements were made for 28 ring structures imaged in 3 separate experiments. Black line – mean MFI; gray border – SD. **D.** Time lapse imaging of GFP-Atg32 in *atg11Δatg32Δ* cells expressing Atg11Δ731-765. Cells were imaged at 15 second intervals, starting 1hr after transfer to SD-N.

To further explore the effects of the CC3 dimerization interface on Atg32 assembly, we examined the shape and brightness of the Atg32 puncta formed in the presence of the different CC3 mutants. In cells expressing Atg11Δ731-765 or Atg11Φ6A the majority of the puncta contained either crescent or ring-shaped concentrations of Atg32 (Figure 4C). These crescent and ring-shaped structures were never observed in cells expressing full-length Atg11. The ring-shaped structures formed by Atg11Δ731-765 were on average as bright as Atg32 puncta formed in the presence of full-length Atg11, but they exhibited a greater range of fluorescence. Their diameter was also significantly greater (0.69 ± 0.22 μm) than standard Atg32 puncta, with a distinct dip in fluorescence in the center (Figure 4D). Time-lapse imaging of GFP-Atg32 in cells expressing Atg11Δ731-765 revealed that Atg32 was initially concentrated into smaller puncta, which then expanded to form the ring shape (Figure 4E). These structures may represent the attempted clustering of Atg32 upon Atg11 binding, suggesting that without the CC3 dimerization interface, proper organization of Atg32 cannot be achieved.

Cells expressing the Atg11-2Ser mutant also had fewer Atg32 puncta than cells expressing full-length Atg11, (Fig. 4A and B), but Atg32 did not form crescent or ring-shaped structures. Instead, all Atg32 puncta formed in the Atg11-2Ser were similar in shape to the Atg32 puncta formed in cells expressing wildtype Atg11. Moreover, we observed GFP in the vacuole of some cells, suggesting that mitophagy was not entirely abrogated.

### The dimerization interface of the Atg11CC3 is required for mitophagy

To investigate whether the CC3 was required for functional mitophagy, we performed the well-established OM45-GFP assay, which utilizes the degradation of the OMM protein OM45 as a readout for mitophagy completion (Kanki et al., 2009a). The OM45-GFP assay was performed in *atg11Δ* cells with different Atg11 variants under the control of their endogenous promoter (Figure 5A and B). As expected, mitophagy was entirely lost in cells lacking Atg11, and expression of full-length Atg11 fully restored mitophagy. In contrast, cells expressing Atg11Δ699-860 or Atg11Δ699-800 had essentially no detectable mitophagy. Moreover, expression of Atg11Δ731- 765 or Atg11Φ6A only supported a 10% restoration of mitophagy compared to full-length Atg11. These data confirm that the dimerization interface of the CC3 is crucial to the proper formation of mitophagy initiation sites and the delivery of mitochondrial cargo to the vacuole. Expression of Atg11-2Ser produced a 64% restoration of mitophagy, supporting the notion that these residues are required for efficient Atg11 function, but that they are less important for mitophagy than the dimerization interface of the CC3. Importantly, there was no significant difference in the expression of the different Atg11 variants, indicating that the reduction in mitophagy levels and the changes in the shape of mitophagy initiation sites are not due to different levels of Atg11 expression (Supplemental Figure S3). We therefore concluded that the Atg11CC3 dimerization interface is required for successful capture of mitochondria during selective autophagy.

**Figure 5.**
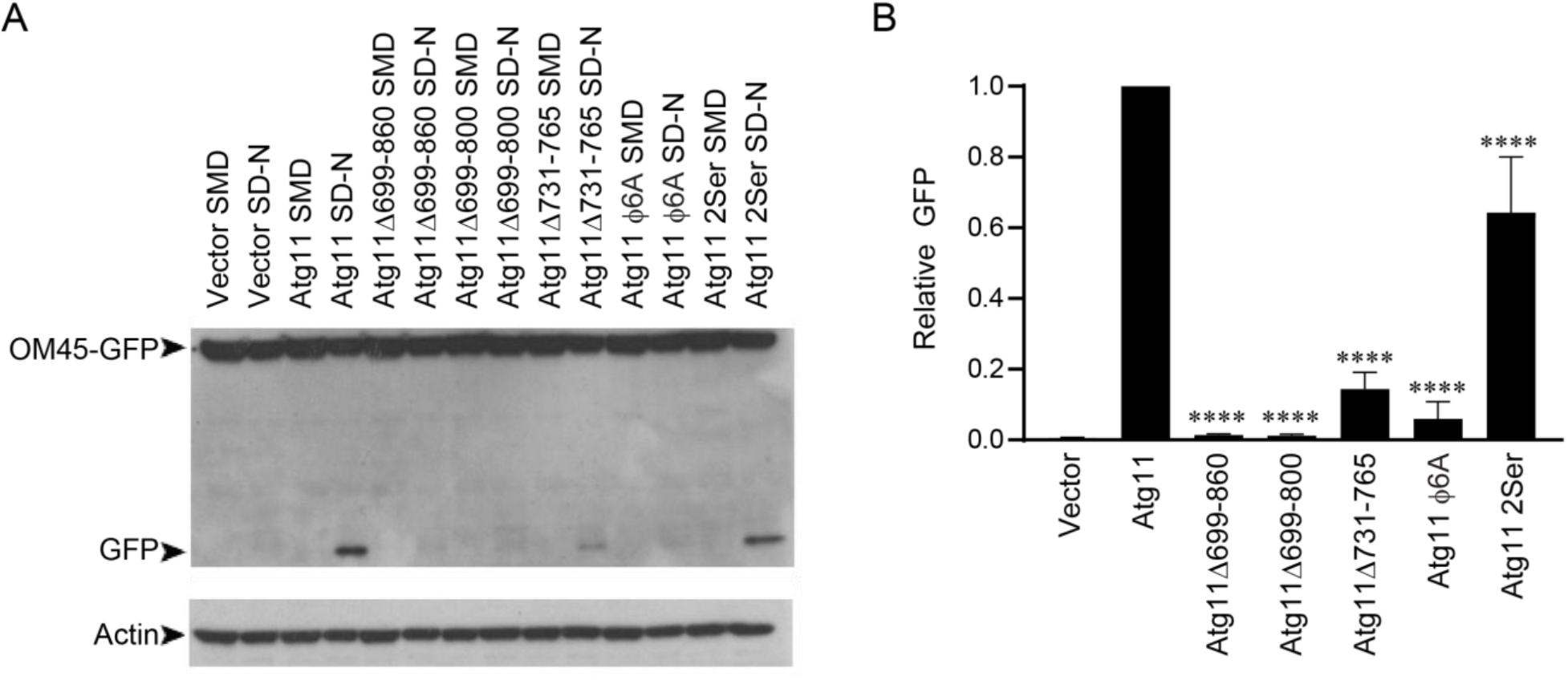
Coiled coil dimer interface mutants in the Atg11CC3 are required for mitophagy. **A.** Atg11 variants were expressed using the endogenous Atg11 promoter in *atg11Δ* cells expressing OM45-GFP. Cells were incubated in SML and transferred to SD-N for 6 hrs and blotted for GFP. An actin loading control is shown below. **B.** Quantification of the GFP bands in each of the SD-N lanes in F. GFP levels were normalized to actin levels, and then to Atg11. Significance was determined using a one-way ANOVA with Sidak’s multiple comparison test, compared to cells expressing Atg11. **** p<0.0001

## Discussion

In this study, we explored the structure and function of the Atg11CC3, a coiled coil domain in Atg11 whose function is not yet known. We show here that the Atg11CC3 forms a parallel coiled coil dimer, and that the dimer interface is crucial for the reorganization of Atg32 into functional mitophagy initiation sites. Taken together our data demonstrates that the CC3 plays an important structural role in mitophagy initiation.

The crystal structure of Atg11_699-800_ revealed that this region is a parallel coiled coil dimer containing a short dimerization interface (residues 730-760) that is flanked on either side by regions containing poor electron density and high B-factors. This may be due to a lack of crystal contacts in these regions or it could suggest that these portions of the CC3 are more dynamic than the dimer interface. If these regions are indeed flexible in full-length Atg11 this may be due to a breakdown of the heptad repeat sequence for residues 724-730 and 766-772. Neither of these stretches contain any hydrophobic residues in the standard positions that would support hydrophobic packing in the center of the helical bundle. If these flanking regions are indeed dynamic in the full-length protein, their flexibility may permit a greater range of movement for the N and C terminal regions of Atg11.

Atg11 has previously been shown to self-associate in cells, and this capacity was primarily assigned to the N-terminal region (Matscheko et al., 2019; Yorimitsu and Klionsky, 2005). However, it was not known how the CC3 contributed to Atg11 self-association. Later studies showed that recombinantly expressed full length Atg11, the N-terminal region of Atg11 and the C-terminal region of Atg11 including the CC3 can all form dimers, making it unclear how each region of Atg11 contributes to the overall self-association of Atg11 in cells (Suzuki and Noda, 2018). It has also been shown that Atg11 may exist in cells as a monomer that dimerizes only once bound to Atg32 (Matscheko et al., 2019). In this study, we found that full length Atg11 is self-associated *in vivo*, independent of nutrient availability. However, we found that the CC3 was entirely dispensable for Atg11 self-association overall. This suggests that Atg11 dimerization is supported primarily by a different interface. Recent work on FIP200, the mammalian analog of Atg11, demonstrated that the N-terminal region of FIP200 is a C shaped dimeric structure (Shi et al., 2020). If Atg11 contains a similar N-terminal region it may serve as the primary site of Atg11 dimerization.

Our data suggests that the CC3 is required for the organization of mitophagy initiation sites. One possible function of the CC3 in the formation of these sites could be as a spacer separating the cargo and the core autophagy machinery. However, we found that deletion of only the CC3 dimer interface (Atg11Δ730-760) or point mutations disrupting the hydrophobic interactions of the dimer interface (Atg11Φ6A) significantly impaired Atg32 puncta formation and almost entirely abolished delivery of mitochondria to the vacuole. Since the Atg11Φ6A mutant contains the same number of amino acids as the wildtype protein this suggests that the function of the CC3 is unlikely to be a spacer.

Previous studies have shown that, in yeast, autophagosomes and mitochondrial fragments captured in autophagosomes are approximately 400 nm in diameter (Backues et al., 2014; Kiššova et al., 2007; Xie et al., 2009). The Atg32 puncta formed in the presence of full length Atg11 are consistent with these measurements. The Atg11 mutants Atg11Δ730-760 and Atg11Φ6A were still able to partially concentrate Atg32 into puncta, but they restructured Atg32 into extended crescent and ring-shaped structures that far exceeded the standard size of autophagosomes and mitochondrial fragments. Tellingly, these structures were not able to progress further in selective autophagy. This suggests that simply concentrating Atg32 is not sufficient to allow for selective autophagy initiation. It seems that Atg32 must also be organized in a specific manner as a necessary precondition for functional mitophagy. We therefore propose that the CC3 plays an important structural role, helping the concentrated Atg32 in the OMM to assume a shape and size suitable for autophagic capture.

A similar principle may hold true for other SARs and their cargos. In yeast, the SAR Atg19 restricts the size of its cargo, the self-associating vacuole protease APE1 (Yamasaki et al., 2016). In mammalian systems, the SAR p62 self-associates, but FIP200 limits cargo size by binding to p62, thereby limiting the size of p62 aggregates (Ciuffa et al., 2015; Jiang et al., 2020). The mammalian ER membrane anchored SAR FAM134B also clusters during selective autophagy, although it is not yet known how the size of FAM134B puncta is regulated (Jiang et al., 2020). Our study suggests that, in the case of yeast mitophagy, Atg11 is required for both concentration of the SAR and restriction of the cargo to a proper size. Thus, the capacity to self-associate may reside in the cargo itself, the SAR, or the adaptor, but in any case, must be regulated to produce cargo suitable for autophagic capture.

Finally, it is important to consider how Atg32 and Atg11 clustering might be influenced by other components of the core autophagy machinery. A recent study demonstrated that the initiation site for non-selective autophagy is formed by liquid-liquid phase separation of the core autophagy machinery, including Atg1, Atg13, Atg17, Atg29, and Atg31 (Fujioka et al., 2020). This phase separation is supported by interactions between Atg13 and Atg17. Atg11 and Atg17 are thought to play counterpart roles in the initiation of selective and non-selective autophagy, respectively. Thus, the proper multimerization of Atg32 and Atg11 may be important for the recruitment and assembly of other elements in the core autophagy machinery, adding another layer of complexity to the mechanism by which mitophagy initiation sites are formed.

## Materials and Methods

### Plasmids

For protein purification, Atg11_699-800_ from S. cerevisiae was codon optimized for *E. coli* and cloned into pHis2 such that the construct contained an N-terminal hexahistidine tag followed by a TEV cleavage site. For all other constructs, Atg32 and Atg11 were amplified from a S. cerevisiae genomic library (Novagen). Atg11 containing 300 base pairs flanking the start and end of the coding region was cloned into yCPLAC33 (Gietz and Sugino, 1988). 2GFP from LD231, which was a gift from Zhiping Xie (Addgene), was then cloned at the N-terminus of Atg11 using Gibson Cloning (NEB) to generate 2GFP-Atg11 yCPLAC33 (Li et al., 2015). sfGFP from the plasmid 1GFP, which was a gift from Scott Gradia, and mCherry were cloned into pCu415CUP1 and pCu416CUP1 vectors, respectively (Labbé and Thiele, 1999). Atg32 was then cloned into GFP pCu415CUP1 and mCherrry pCu416CUP1. HA or FLAG tags were added to Atg11 using PCR and these were cloned into pCu415CUP1 and pCu416CUP1, respectively. Point mutants and truncations were generated using Q5 Site-Directed Mutagenesis (NEB) and all mutagenesis primers were designed using NEBaseChanger (http://nebasechanger.neb.com/).

### Yeast Strains

*atg1Δ* and *atg11Δ* cell lines were obtained from the S. cerevisiae knockout collection (Invitrogen). atg32 was replaced with the NAT cassette from pAG25 (Goldstein and McCusker, 1999) in the *atg1Δ* and *atg11Δ* cell lines to generate KBY006 (MATα *his3Δ1 leu2Δ0 lys2Δ0 ura3Δ0 atg1Δ::KAN atg32Δ::NAT*) and SKY004 (MATα *his3Δ1 leu2Δ0 lys2Δ0 ura3Δ0 atg11Δ::KAN atg32Δ::NAT*). TKYM22 (SEY6210 *OM45-GFP::TRP1*) was a gift from Daniel Klionsky (Kanki and Klionsky, 2008; Mao et al., 2011). atg11 was replaced with the NAT cassette from pAG25 in TKYM22 to generate XXY003 (TKYM22 *atg11Δ::NAT*).

### Microscopy

Cells were grown in SMD (0.67% yeast nitrogen base, 2% glucose and supplemented with the appropriate amino acid and vitamin mixture) with 0.5uM copper unless otherwise stated, to an OD of 1. For SD-N treatment, cells were washed once in SD-N (0.17% yeast nitrogen base without amino acids and ammonium sulfate, 2% glucose), then resuspended in SD-N and incubated for 1 hr. To visualize mitochondria, cells were stained with 50nM Mitotracker red CMXRos (Invitrogen) for 10 minutes, then washed in the appropriate media (SMD or SD-N) and imaged.

Cells were plated in SMD or SD-N media on MatTek glass bottom dishes and allowed to settle for 5 minutes before imaging. Images were acquired on a Zeiss Airyscan LSM 880 microscope using an alpha Plan-Apochromatic 100x oil lens with NA=1.46 using Zeiss Immersol 518 F immersion oil at room temperature. GFP was excited using a 488 Argon laser, mCherry and CMXRos Mitotracker were excited using a DPSS 561 laser. Emission was detected using a Zeiss GaAsP-PMT detector. Cells were imaged in z-stacks to a final height of 6um. For time lapse imaging, cells were imaged as described above at 15 second intervals. Acquisition was performed using the ZEN black software and analyzed using ZEN black, ZEN blue and ImageJ software. Airyscan images were deconvoluted by Airyscan processing set to 3D with a processing strength of 6, and the 3D stacks were used to generate 3D max projections. No gamma correction was performed.

Puncta were defined based on the phenotype observed in *atg1Δatg32Δ* cells expressing GFP- Atg32 that were incubated in SD-N for 1 hr (Figure 1B). The experiment was performed in 3 independent repeats, and >50 puncta were analyzed in total. MFI values were obtained from raw data and the fluorescence values measured outside cells was subtracted. The MFI of puncta was normalized to the MFI of mitochondria directly adjacent. For puncta diameter measurements, each trace was analyzed independently.

For puncta quantification, each experiment was performed in 3 independent repeats. In each repeat, at least 6 fields were acquired from different regions of the dish, yielding >300 cells imaged per group per repeat. All the cells imaged were analyzed, totaling >1000 cells per group. Since GFP was entirely absent from the vacuole of *atg1Δatg32Δ* and *atg11Δatg32Δ* cells, we determined that GFP delivery to the vacuole was entirely dependent on selective autophagy as described previously (Xia et al., 2018). We therefore categorized cells with vacuolar GFP as having recently contain GFP-Atg32 puncta.

### Statistical Analysis

Statistical analysis was performed using GraphPad Prism 7.0. Data was compared using one-way ANOVA with Dunnet’s multiple comparison tests (Figure 4G) or two-way ANOVA with Sidak’s multiple comparison tests (Figure 1E and 4B). * p<0.05, *** p<0.001, **** p<0.0001.

### Protein Expression and Purification

Bl21(DE3)-RIL cells transformed with Atg11_699-800_ in pHis2 were grown in LB (Fisher Scientific) at 37°C to an OD600 between 0.6-0.8. Cells were incubated at 4°C while the incubator cooled to 18°C. Expression was then induced with 1 mM IPTG and cells were grown at 18°C overnight. Cells were harvested by centrifugation at 4000 x g for twenty-five minutes at 4°C, then resuspended in 30 mL lysis buffer containing 50 mM Tris pH 8.0, 500 mM NaCl, 0.1% Triton X-100 (Alfa Aesar), 166 μM MgCl, with a cOmplete, EDTA-free Protease Inhibitor tablet (Sigma Aldrich) and benzonase (Sigma Aldrich). Cells were lysed using a French Press (Thermo Electron) and lysates were cleared by centrifugation at 40,000 x g at 4°C. Cleared lysate was subjected to purification using TALON resin (Clontech) which was equilibrated with 50 mM Tris, pH 8.0, 500 mM NaCl and protein was eluted with 20 mM Tris, pH 8.0, 200 mM NaCl, 200 mM Imidazole. The hexahistidine tag was cleaved overnight with recombinant TEV protease at 4°C. Atg11699-800 was further purified with a second TALON resin purification followed by size exclusion chromatography using a HiLoad Superdex 75 PG column (GE Healthcare) equilibrated in 20 mM Tris pH 8.0, 200 mM NaCl.

For selenomethionine labeling, Bl21(DE3)-RIL cells transformed with Atg11_699-800_ in pHis2 were grown in M9 minimal media with 2% glucose at 37°C to an OD600 of 0.5-0.6. Methionine production was blocked using an amino acid mixture containing 50 mg leucine (Alfa Aesar), 50 mg isoleucine (Tokyo Chemical Industry), 50 mg valine (Alfa Aesar), 100 mg lysine (Tokyo Chemical Industry), 100 mg phenylalanine (Tokyo Chemical Industry), 100 mg threonine (Tokyo Chemical Industry) per liter of media. 60 mg of selenomethionine (Acros Organics) was also added to each liter of media. After fifteen minutes of additional growth at 37°C, cells were stored at 4°C while the incubator cooled to 18°C and then protein expression was induced with 1 mM IPTG (VWR) and expressed at 18°C overnight. Protein was purified as above but with 5 mM 2- mercaptoethanol added to all lysis and talon buffers and with 0.2 mM tris(2- carboxyethyl)phosphine added to the size exclusion buffer.

### Crystallization and Structure Determination of Atg11_699-800_

Atg11_699-800_ was concentrated to 11.8 mg/mL using Amicon ultra 3000 concentrators (Millipore). After optimization, rod-shaped crystals containing native protein grew within one day at room temperature in 90 mM LiNaK, 0.1 M Morpheus II buffer system 4 pH 6.5, 50% v/v Morpheus II precipitant mix 6 using a protein to reservoir ratio of 1:2. Crystals containing selenomethionine labeled protein grew in similar conditions (90 mM LiNaK, 0.1 M Morpheus buffer system 4 pH 6.5, 10.5% PEG 4k, 15% 1,2,6 hexanetriol protein to reservoir ratio 1:1). Anomalous data on selenomethionine crystals were collected at NE-CAT (24-ID-C) at APS while data was collected on native protein crystals at FMX (17-ID-2) at NSLSII. Images were processed in XDS (Kabsch, 2010) and scaled using AIMLESS (Evans and Murshudov, 2013). Autosol (Terwilliger et al., 2009) within Phenix (Liebschner et al., 2019) was used for SAD phasing. An initial model was built in Coot (Emsley et al., 2010)and this partial model was then used as an MR search model in Phenix with the higher resolution native data set. Subsequently, iterative model building in Coot and refinement in Refmac (Murshudov et al., 1997) in CCP4 (Winn et al., 2011) were performed. The final structure was deposited in the PDB as 6VZF.

### Co-Immunoprecipitation

*atg11Δ* or SKY004 cells were transformed with the appropriate vectors. Cells were grown to mid-log phase in SMD. Protein expression was then induced with 50 μM copper (Sigma). After 3 hrs of induction, cells were pelleted (2,000 x g, 5 min) and either stored at −80°C for SMD conditions or resuspended in SD-N with 1 mM PMSF (Amresco) for 30 min and then harvested and stored at −80°C. Cells were resuspended in lysis buffer containing 1X phosphate-buffered saline (Corning), 0.2 M sorbitol (Sigma), 1 mM MgCl2 (VWR), 1% Triton X-100 (Alfa Aesar), 1 mM PMSF, and 1 cOmplete EDTA-free tablet (Sigma Aldrich). Cells were lysed by bead beating and centrifuged at 16,100 x g at 4°C. Supernatant was incubated with FLAG M2 magnetic beads (Sigma), rocking for 2 hrs at 4°C. After removal of the flow-through, beads were washed 5 times with lysis buffer. 1 X NuPAGE LDS sample buffer (Invitrogen) was added to each tube. Tubes were then incubated at 37°C for 30 min to elute proteins bound to the magnetic beads. Supernatant was analyzed by western blot and probed with anti-FLAG (Sigma), anti-HA (Abcam) or anti-GFP (Santa Cruz) antibodies.

### OM45-GFP Processing Assay

The OM45-GFP processing assay was performed as previously described (Kanki et al., 2009a). XXY003 cells were transformed with the appropriate vectors to add back atg11 variants. Cells were grown at 30oC in SMD to an OD600 of 0.9-1.0, pelleted at 1,400 x g for 10 min and resuspended in SML (0.67% yeast nitrogen base, 2% lactic acid and supplemented with the appropriate amino acid and vitamin mixture). After growing to an OD600 of 0.8-1.5, cells were pelleted and resuspended cells in SD-N for six hours. After incubation in SD-N, cells were harvested, lysed by bead beating and GFP was detected by western blot.

## Acknowledgements

This work was supported by R35GM128663 and P20GM113132. We would like to thank Dan Klionsky for reagents. DNA sequencing was performed using the Molecular Biology Shared Resource at Dartmouth which is supported by P30CA023108. Crystallography data were collected at NE-CAT (24-ID-C) at APS and FMX (17-ID-2) at NSLSII. This work is based upon research conducted at the Northeastern Collaborative Access Team beamlines, which are funded by the National Institute of General Medical Sciences from the National Institutes of Health (P30 GM124165). The Pilatus 6M detector on 24-ID-C beam line is funded by a NIH-ORIP HEI grant (S10 RR029205). This research used resources of the Advanced Photon Source, a U.S. Department of Energy (DOE) Office of Science User Facility operated for the DOE Office of Science by Argonne National Laboratory under Contract No. DE-AC02-06CH11357. The Center for Biomolecular Structure (CBMS) operation of the FMX beam line at NSLS II is primarily supported by the National Institute of Health, National Institute of General Medical Sciences (NIGMS) through a Core Center of Excellence P30 Grant, (1P30GM133893) and by the DOE Office of Biological and Environmental Research (BER) through BER-BO 070 grant. As a National facility resource, the National Synchrotron Light Source II at Brookhaven National Laboratory, is supported by the U.S. Department of Energy (DOE). Worked performed at the CBMS facilities are supported in part by the DOE Office of Science, Office of Basic Energy Sciences Program (BES), BES-FWP-PS001.

