## Supplemental Materials for "The third coiled coil domain of Atg11 is required for shaping mitophagy initiation sites"

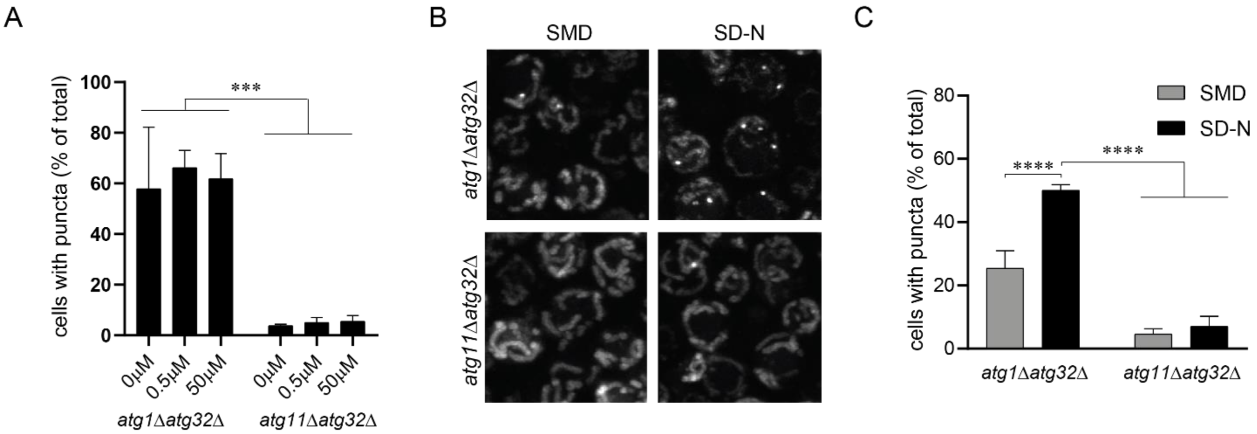


**Figure S1.** The formation of Atg32 puncta is independent of Atg32 concentration and fluorophore. **A.** Quantification of puncta prevalence observed in atg1Δatg32Δ or atg11Δatg32Δ cells expressing GFP-Atg32 driven by the CUP1 promoter, induced with the indicated copper concentrations and incubated in SD-N for 1 hr. **B.** Representative images showing mCherry-Atg32 driven by the CUP1 promoter with 0.5 μM copper in atg1Δatg32Δ or atg11Δatg32Δ cells. Cells were grown in SMD, then transferred to SD-N for 1 hr. **C.** Quantification of the percent of cells containing puncta from B.

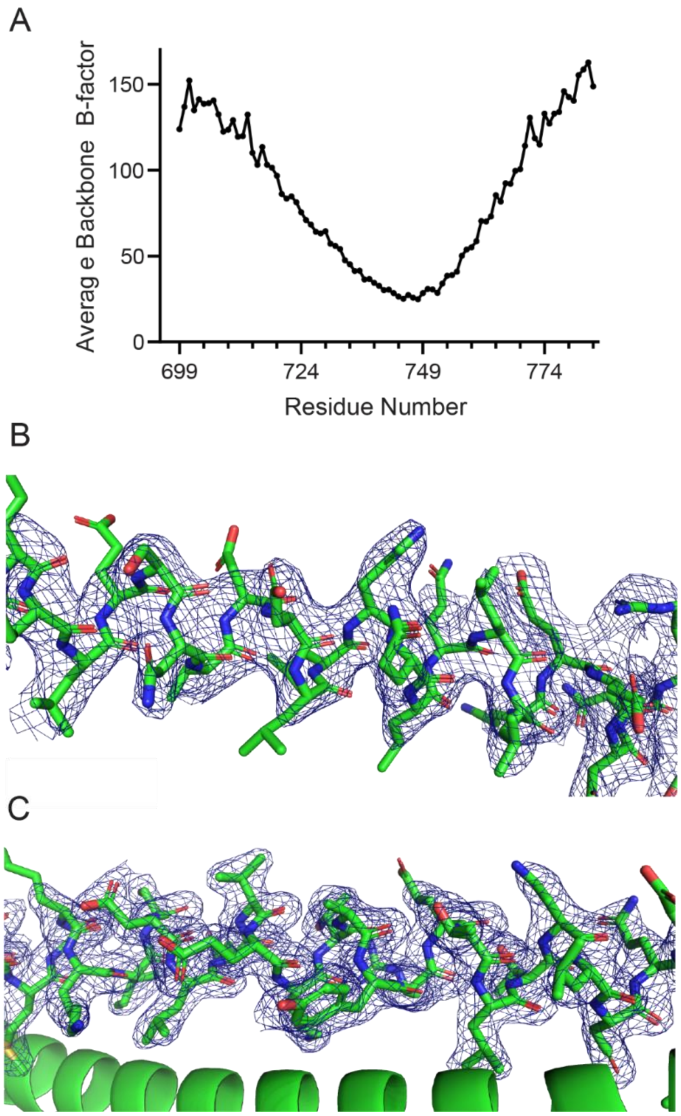


**Figure S2.** B-factors and Electron Density for Atg11$_{699\text{-}800}$. **A.** Average B-factor values for backbone atoms determined by BAVERAGE in CCP4 and plotted per residue. **B.** 2Fo-Fc map contoured at 1.0 σ for residues 704-726. **C.** 2Fo-Fc map contoured at 2.0 σ for residues 734-757.

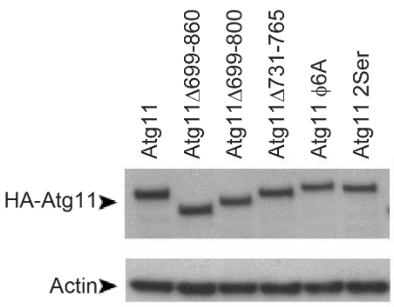


**Figure S3.** Expression of different Atg11 variants. HA-Atg11 variants were expressed in *atg11Δatg32Δ* cells grown in SMD. Samples were blotted against HA.
